# The glycolytic reaction PGAM unexpectedly restrains Th17 pathogenicity and Th17-dependent autoimmunity

**DOI:** 10.1101/2024.08.18.607992

**Authors:** Chao Wang, Allon Wagner, Johannes Fessler, David DeTomaso, Sarah Zaghouani, Yulin Zhou, Kerry Pierce, Raymond A. Sobel, Clary Clish, Nir Yosef, Vijay K. Kuchroo

## Abstract

Glucose metabolism is a critical regulator of T cell function, largely thought to support their activation and effector differentiation. Here, we investigate the relevance of individual glycolytic reactions in determining the pathogenicity of T helper 17 (Th17) cells using single-cell RNA-seq and Compass, an algorithm we previously developed for estimating metabolic flux from single-cell transcriptomes. Surprisingly, Compass predicted that the metabolic shunt between 3-phosphoglycerate (3PG) and 2-phosphoglycerate (2PG) is inversely correlated with pathogenicity in these cells, whereas both its upstream and downstream reactions were positively correlated. Perturbation of phosphoglycerate mutase (PGAM), an enzyme required for 3PG to 2PG conversion, resulted in an increase in protein expression of IL2, IL17, and TNFa, as well as induction of a pathogenic gene expression program. Consistent with PGAM playing a pro-regulatory role, inhibiting PGAM in Th17 cells resulted in exacerbated autoimmune responses in the adoptive transfer model of experimental autoimmune encephalomyelitis (EAE). Finally, we further investigated the effects of modulating glucose concentration on Th17 cells in culture. Th17 cells differentiated under high- and low-glucose conditions substantially differed in their metabolic and effector transcriptomic programs, both central to Th17 function. Importantly, the PGAM-dependent gene module marks the least pathogenic state of Th17 cells irrespective of glucose concentration. Overall, our study identifies PGAM, contrary to other glycolytic enzymes, as a negative regulator of Th17 pathogenicity.

## INTRODUCTION

Glycolysis is central to T helper 17 (Th17) cell differentiation and function. Genetic and pharmacological inhibition of glycolytic and related enzymes, such as HIF1a^1^, PKM2^2^, GLUT3^3^, PDHK^4^, LDHA^5^, and GPI1^6^, have all resulted in the loss of Th17 cell differentiation or effector functions. These studies, as well as extensive literature on other immune cells, have led to the concept that glycolysis promotes pro-inflammatory functions in immune cells^7–13^. However, the role of glycolysis in the induction of pro-inflammatory phenotypes may be more nuanced^14,15^. For example, inhibition of glycolysis with 2-deoxy-glucose (2DG) promotes Th17 cell differentiation^16^. Glycolysis is also necessary for thymus-derived regulatory T cell proliferation and functional fitness^15^. These observations suggest that suppression of glycolysis is not a general anti-inflammatory strategy. Similarly, other metabolic pathways, such as fatty acid oxidation, can either support anti-inflammatory responses or promote inflammasome activation in immune cells^14^. We thus decided to study the connection between individual reactions along the glycolysis pathway and the function of Th17 cells.

In this study, we focus on the Th17 subset of T helper cells, leveraging single-cell RNA sequencing (scRNA-seq) and Compass^17^, an algorithm we developed and validated for in silico modeling of metabolic states of single cells. Th17 cells demonstrate heterogeneous effector profiles and functions in vivo^18^. Distinct subtypes of Th17 cells can be obtained in vitro, depending on the culture conditions used to differentiate them. Th17 cells differentiated under pathogenic conditions (Th17p) induce severe autoimmune encephalomyelitis (EAE) upon adoptive transfer. Conversely, transfer of Th17 cells derived under non-pathogenic conditions (Th17n) results in mild to no EAE^19–21^. We tested the correlation between the activity of glycolytic reactions (predicted with Compass) and the functional state of Th17 cells (estimated using a signature of pro-inflammatory and pro-regulatory transcriptional modules we have previously identified^22^). We identified several glycolytic enzymes that are significantly associated with the pathogenicity of Th17 cells either positively or, unexpectedly, also negatively. Perturbation of G6PD and PGAM in Th17 cells confirmed these predicted opposing functions; G6PD inhibition restricted the pathogenic gene expression module, whereas PGAM inhibition shifted the cells towards a pathogenic state and exacerbated autoimmunity and tissue inflammation in the EAE murine model. We then investigated the impact of glucose availability on the Th17 effector phenotype and discovered unique transcriptional responses associated with glucose scarcity or abundance. Th17p cells manifested two distinct effector programs, a CSF2+GZMB+ program under low-glucose condition versus a TBX21+ program under high-glucose conditions. In Th17n cells, a pro-regulatory program was detected under both low- and high-glucose conditions, but was considerably restricted in the former. PGAM inhibition specifically targeted this pro-regulatory program, leading to the induction of a pro-inflammatory state in Th17n cells.

## RESULTS

### The glycolytic enzyme phosphoglycerate mutase (PGAM) suppresses Th17 cell pathogenicity

To analyze the effects of glycolytic reactions on the function of T helper cells, we analyzed scRNA-seq data from 1,311 Th17n cells derived from naïve CD4+ T cells activated and differentiated in vitro under non-pathogenic Th17 culture conditions (TGFb1+IL6). Although cells differentiated in this setting do not evoke strong inflammation in the EAE model, we and others have demonstrated that Th17n cells actually span a spectrum of functional states, expressing different proportions of pro-inflammatory and pro-regulatory cytokines and transcription factors. Driving Th17n cells to the pathogenic limit of this spectrum through perturbation of key regulators can indeed result in inflammation^20,22^. We hypothesized that the more pro-inflammatory cells in the Th17n spectrum could be more glycolytic than the pro-regulatory cells. To test this, we defined a computational pathogenicity score for Th17n cells based on our previously discovered Th17n pro-inflammatory and pro-regulatory transcriptional modules^22^. We define the pathogenicity score of each cell as the difference between the sum of normalized expression of 63 pro-inflammatory genes and the sum of 30 pro-regulatory genes **(Table S1; Methods)**.

We next applied Compass to the scRNA-seq data to approximate the activity of ∼900 metabolic reactions in each cell, including dozens of reactions associated with central carbon metabolism, of which 19 were associated with the glycolysis pathway. For each reaction, we computed the Spearman correlation between the Compass activity scores and the pathogenicity scores across all cells, and ranked the reactions by their correlation with Th17n pathogenicity **(Figure 1A-B)**. The results surprisingly suggested that the glycolytic pathway does not behave as a cohesive unit. Instead, metabolic reactions between 3-phosphoglycerate (3PG) and phosphoenolpyruvate (PEP) were negatively correlated with the pathogenicity score, namely positively associated with a pro-regulatory Th17n phenotype. Compass further predicted that the metabolic shunt from 3-phosphoglycerate (3PG) towards serine biosynthesis was also negatively correlated with the pathogenicity score. This prediction suggests that carbon fate between 3PG and PEP of the glycolysis pathway may be important for the pro-regulatory Th17n effector program, contrary to the prior expectation that higher glycolytic activity would always be associated with the pro-inflammatory Th17 program. Other major glycolytic reactions, including Lactate dehydrogenase (LDH), pyruvate dehydrogenase (PDH), and phosphoenolpyruvate carboxykinase (PCK) were all positively correlated with the pathogenicity score, namely a pro-inflammatory Th17 phenotype, suggesting that carbon derived from precursors other than 3PG may be used to obtain pyruvate in pro-inflammatory Th17n cells.

**Figure 1.**
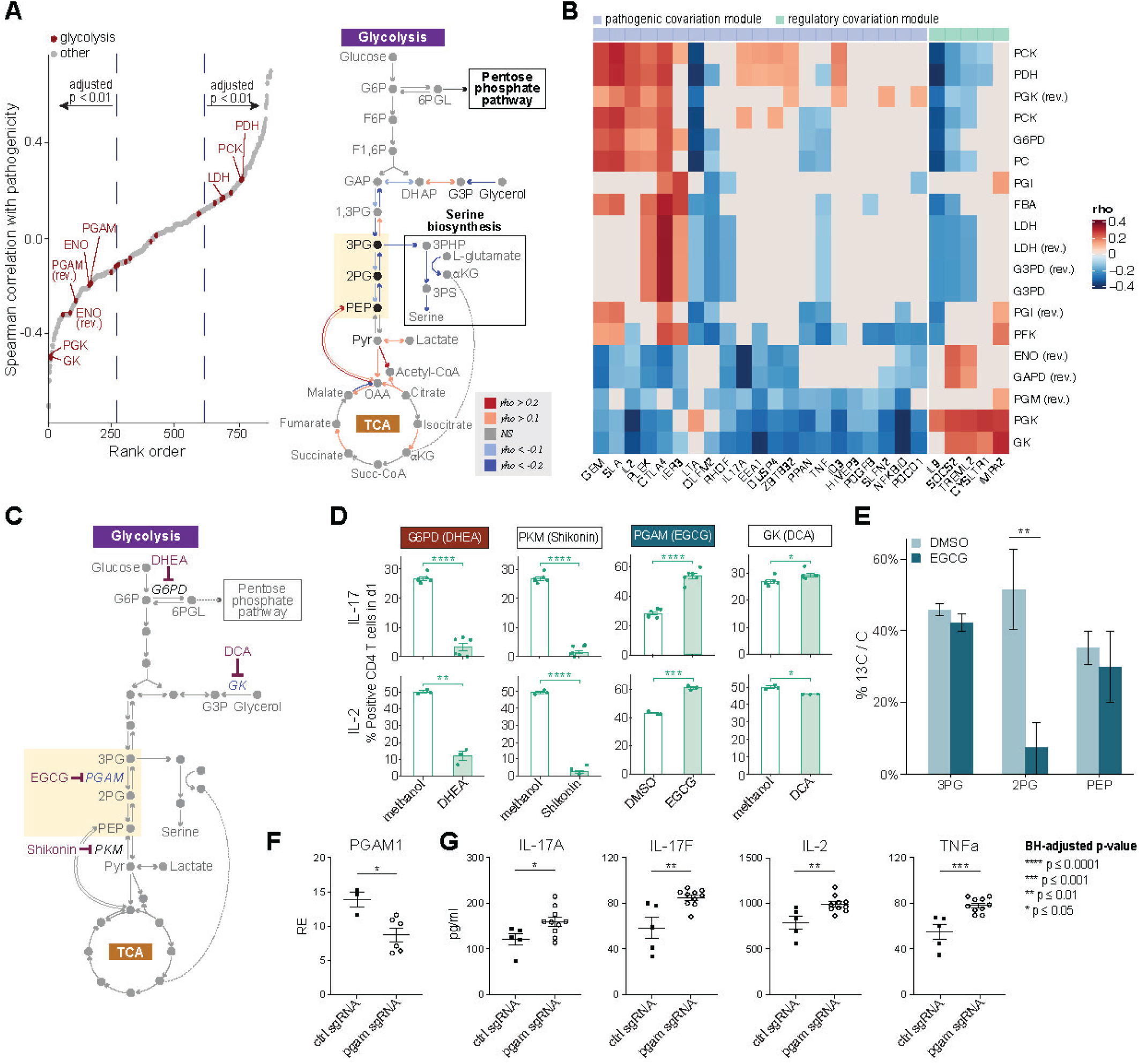
The glycolytic enzyme phosphoglycerate mutase (PGAM) suppresses Th17 cell pathogenicity. (A) Left: Metabolic reactions were ordered by the Spearman correlation of their Compass-predicted activity with a Th17 pathogenicity signature based on previously inferred transcriptomic modules **(Methods)**. Right: A schematic diagram of central carbon metabolism. Reactions are colored according to the Spearman correlation of their Compass-predicted activity with the Th17 pathogenicity signature (reactions with a non-significant correlation shown in grey). (B) Spearman correlations between Compass-predicted activity of glycolysis reactions (rows) and genes belonging to the pro-inflammatory or pro-regulatory transcriptomic modules. Only statistically significant correlations with |rho| > 0.25 are shown in color; only reactions/genes with at least one colored entry are shown. (C) Inhibitors of selected reactions in central carbon metabolism are denoted. (D) Th17 cytokines were measured by flow cytometry. Naïve T cells were differentiated under non-pathogenic Th17 (Th17n) conditions **(Methods)** in the presence of control solvent or inhibitors. Cells were pre-labeled with division dye; protein expression is reported for cells that have gone through one division (d1) to exclude arrested cells. (E) ratio of 13C-tagged carbon to total carbon in metabolites retrieved from Th17 cells cultured for 15 minutes in the presence of 13C-glucose. Three metabolites are shown: PGAM’s substrate (3-phosphoglycerate), product (2-phosphoglycerate), and the next downstream metabolite along the glycolytic pathway (phosphoenolpyruvate). (F) RT-qPCR results showing “knockdown” of PGAM in Th17n cells where each dot represents an independent lentiviral infection with either control or PGAM sgRNA containing lentiviruses. (E-F) Statistical significance computed by Student’s t-test with a BH adjustment for multiple comparisons where appropriate. Reaction and metabolite abbreviations appear in Table XXX.

To validate whether metabolic reactions between G3P and PEP were indeed associated with the pro-regulatory Th17n phenotype, we chemically inhibited the enzymes that catalyze the upstream segment (glucose-6-phosphate dehydrogenase [G6PD]), the segment itself (phosphoglycerate mutase [PGAM]), and the downstream segment (pyruvate kinase muscle isozyme [PKM]) (**Figure 1C)**. We also inhibited the enzyme catalyzing the reaction glycerol kinase (GK), also predicted to be negatively correlated with pathogenicity **(Figure 1A)**. The inhibitors were dehydroepiandrosterone (DHEA, inhibits G6PD), epigallocatechin-3-gallate (EGCG, inhibits PGAM), and shikonin (inhibits PKM2), and 2,3-dihydroxypropyl-dichloroacetate (DCA, inhibits GK) (**Methods**).

We first analyzed the effects of these inhibitors on the differentiation and function of Th17n cells using flow cytometry (**Figure 1D**). Due to the possibly deleterious effects of blocking these central reactions, we used the highest dose of each inhibitor that did not affect cell viability (compared to solvent alone). We further used flow cytometry to restrict the analysis to cells that had undergone one division (d1), thereby excluding arrested cells and cells that have been blocked from activation and expansion. In addition, each treatment group was matched with an appropriate vehicle control (**Figure 1D**). We found that the percentages of IL17 and IL2 expressing Th17n cells were significantly upregulated by chemical inhibition of the glycolytic PGAM, in agreement with the prediction made by Compass. IL17 expression also slightly increased following the chemical inhibition of GK, which was also predicted to be negatively associated with the pathogenic phenotype, but significantly decreased following chemical inhibition of the G6PD and PKM reactions, that were predicted to be positively associated with the pathogenic Th17 module **(Figure 1D)**. We then quantified the secretion of a larger set of Th17 cytokines and found that inhibition of PKM or G6PD curtailed all cytokine production, suggesting that these enzymes are important for overall T cell effector functions. In contrast, cells subject to PGAM or PKM inhibition retained their cytokine profile with a few exceptions **(Figure S1A).**

To verify that the effect of EGCG was mediated by inhibition of PGAM we conducted a carbon tracing assay in which the cell’s medium was supplemented with 13C-glucose **(Methods)**. PGAM inhibition with EGCG led to a sharp decrease (from 51% 13C ratio to 7% in PGAM’s immediate product, 2-phosphoglycerate (2PG), but not in the PGAM substrate 3-phosphoglycerate (3PG) or phosphoenolpyruvate (PEP), which is one step downstream of 2PG in **(Figure 1E)**. We did not detect a significant decrease in 13C ratio in any other glycolytic metabolite we measured **(Table S1**). This suggests that the effect of the inhibitor EGCG is restricted (at least within glycolysis) to the PGAM reaction.

Since, chemical inhibtion with EGCG could have off-target effects, we next studied the effect of genetic perturbation of PGAM on Th17 cell function. To this end, we differentiated Th17n cells from naïve CD4 T cells isolated from Cas9 transgenic mice and transfected them with either control or sgRNA against PGAM. We confirmed that PGAM expression is significantly reduced in PGAM sgRNA as compared to control sgRNA transfected cells (**Figure 1F**). We found that genetic deletion of PGAM significantly promoted the secretion of IL17A, IL17F, IL2, and TNFa protein from Th17n cells, largely consistent with the effect of the chemical inhibitor EGCG (**Figure 1G**).

Taken together, Compass analysis predicted that within the Th17n compartment, the glycolytic PGAM reaction inhibits, rather than promotes, the capacity of the cells to induce pathogenic Th17 phenotype. Consistently, PGAM perturbation promoted the expression of proinflammatory cytokines IL17, IL2, and TNFa.

### PGAM inhibition promotes transcriptomic shifts toward a pathogenic state in Th17 cells

To analyze the broader impact of perturbing glycolytic enzymes on the transcriptome, we used bulk RNA-Seq to profile Th17n and Th17p cells grown in the presence of either the predicted pro-regulatory inhibitor DHEA (inhibiting G6PD) or the predicted pro-inflammatory inhibitor EGCG (inhibiting PGAM) and their respective controls (**Figure 2A**).

**Figure 2.**
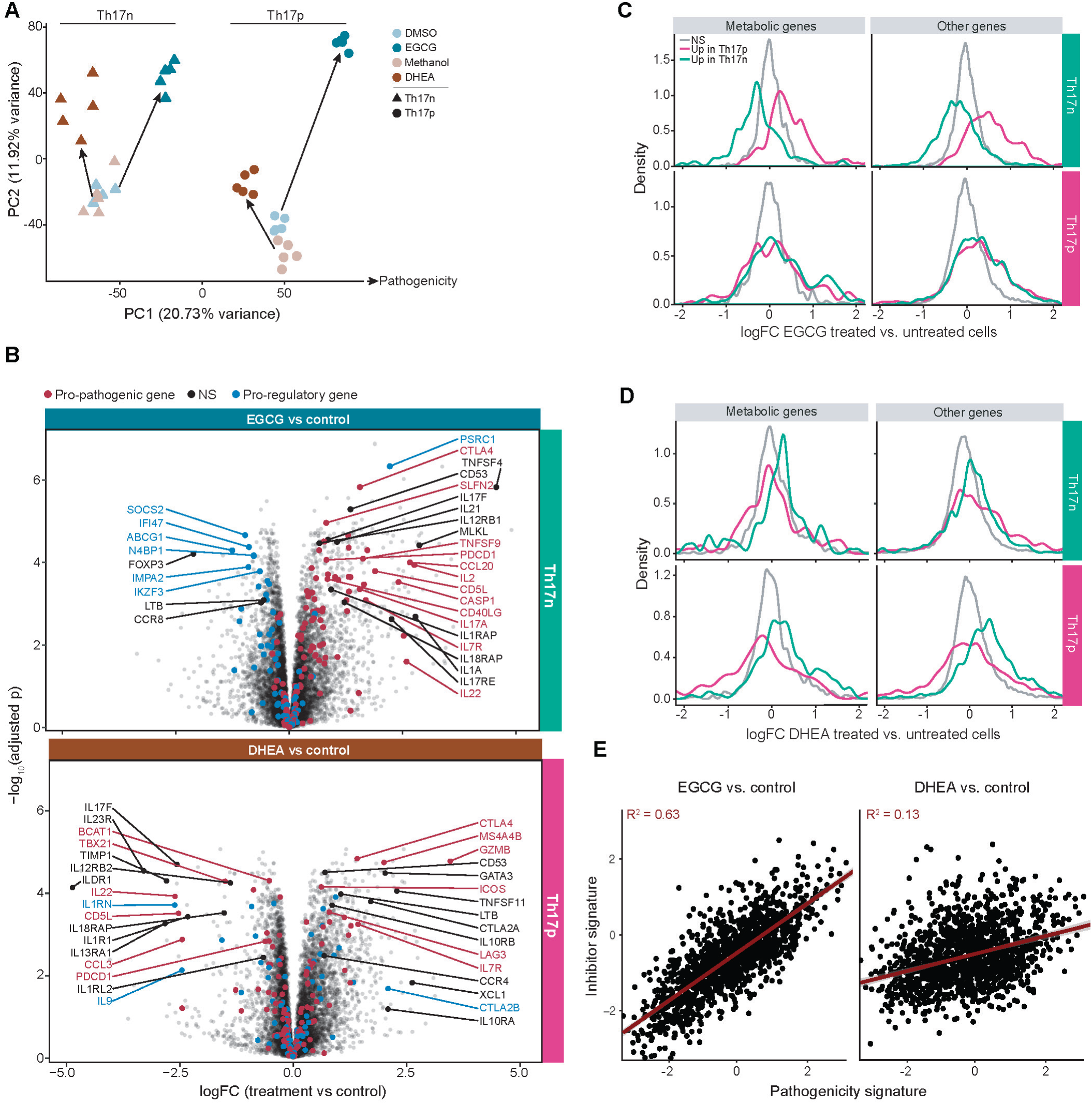
PGAM inhibition promotes transcriptomic shifts toward a pathogenic state in Th17 cells. (A) PCA of bulk RNA-Seq of d1 Th17 cells. (B) Differential gene expression due to EGCG and DHEA treatment. Red and blue dots represent genes associated with the pathogenic and regulatory Th17 states, respectively. Red genes belong to the previously defined pro-pathogenic transcriptomic module^22^ or a list of pathogenic Th17 markers^19^. Blue genes are similarly defined. (C-D) Histograms of the logFC per gene in differential expression of (C) EGCG-treated or (D) DHEA-treated vs. DMSO-treated cells. A separate histogram is shown for Th17p-associated (magenta), Th17n-associated (green), and non-significantly associated (grey) genes. Genes were partitioned into these three groups by differential expression in bulk RNA-Seq (same libraries as shown in panel A) between DMSO-treated Th17p and Th17n cells with a significance threshold of BH-adjusted p ≤ 0.05 and log2 foldchange ≥ 1.5 in absolute value. (G) ratio of 13C-tagged carbon to total carbon in Th17 cells. (E) Two transcriptomic signatures were computed for each Th17n cell (dots): its pathogenicity score (as in Figure 1) and a score reflecting its similarity to EGCG-treated or DHEA-treated cells comparted to DMSO-treated cells **(Methods)**.

Differential expression analysis with the resulting data **(Figure 2B-C; Table S2)** suggests that EGCG strengthened the pathogenic transcriptional program of Th17n cells, upregulating pro-inflammatory genes (e.g., IL22, IL7R, and CASP1) and (to a more limited extent) downregulating pro-regulatory ones (e.g., IKZF3) **(Figure 2B, upper panel)**. DHEA, in contrast, upregulated some pro-inflammatory genes (e.g., GZMB) but suppressed others (e.g., TBX21, IL22) **(Figure 2B, lower panel)**. To further probe how systematic this effect may be, we divided the transcriptome into three groups – genes that are preferentially expressed by Th17p cells (magenta curve), by Th17n cells (green curve), or expressed similarly by both (grey curve). We detected a clear upregulation of the pathogenic (magenta) program by EGCG together with downregulation of the non-pathogenic (green) program in Th17n) **(Figure 2C)**. A similar trend was observed when considering genes that encode metabolic enzymes, which supported the hypothesis that inhibition of the metabolic reaction catalyzed by PGAM with EGCG orchestrated a network-wide metabolic shift that resulted in the emergence of a pro-inflammatory Th17 program **(Figure 2C)**. In contrast, DHEA only promoted the pro-regulatory (green) program in Th17p cells **(Figure 2D)**. The already expressed pro-pathogenic program did not.

Lastly, we used the differential expression results to derive a gene signature of PGAM inhibition by EGCG on Th17n cells **(Methods; Table S2)**. We scored each (untreated) Th17n cell in our scRNAseq data according to its expression of this signature, thereby placing it on the spectrum of transcriptional states that span between EGCG and vehicle treatments. We found that this signature, which reflects the effects of EGCG, is strongly correlated (Pearson rho = 0.79; R^2^ = 0.63) with the Th17 pathogenicity signature **(Figure 2E, left)**. A similarly derived gene signature of DHEA effects on Th17n had a weaker association with the pathogenicity score (Pearson rho = 0.36; R^2^ = 0.13) **(Figure 2E, right)**.

### PGAM inhibition suppresses a regulatory Th17n program

To further study the transcriptional effects of EGCG and DHEA, we supplemented our scRNAseq data, which was generated following standard protocols using 25mM glucose in media, with Th17 cells differentiated under glucose-poor media (1mM). We hypothesized, that the G6PD- and PGAM-dependent regulation of Th17 cell pathogenicity may reflect an arrest of glucose utilization. As expected, changes in glucose abundance induced a large shift in Th17n and Th17p gene expression programs **(Figure 3A, Figures S2A)** and promoted cell proliferation **(Figure S2B)**. Furthermore, the lower glucose concentration significantly curtailed cellular metabolic activity (defined as the portion of the transcriptome consisting of RNA coding for metabolic enzymes) in both Th17n and Th17p cells **(Figure 3B).** The pathogenicity score of Th17n was significantly higher in the low-glucose conditions **(Figure 3C)**. The increase can be attributed to a significant mitigation of the pro-regulatory module under glucose-poor conditions, and not to an enhancement of the pro-inflammatory module **(Figure 3C).**

**Figure 3.**
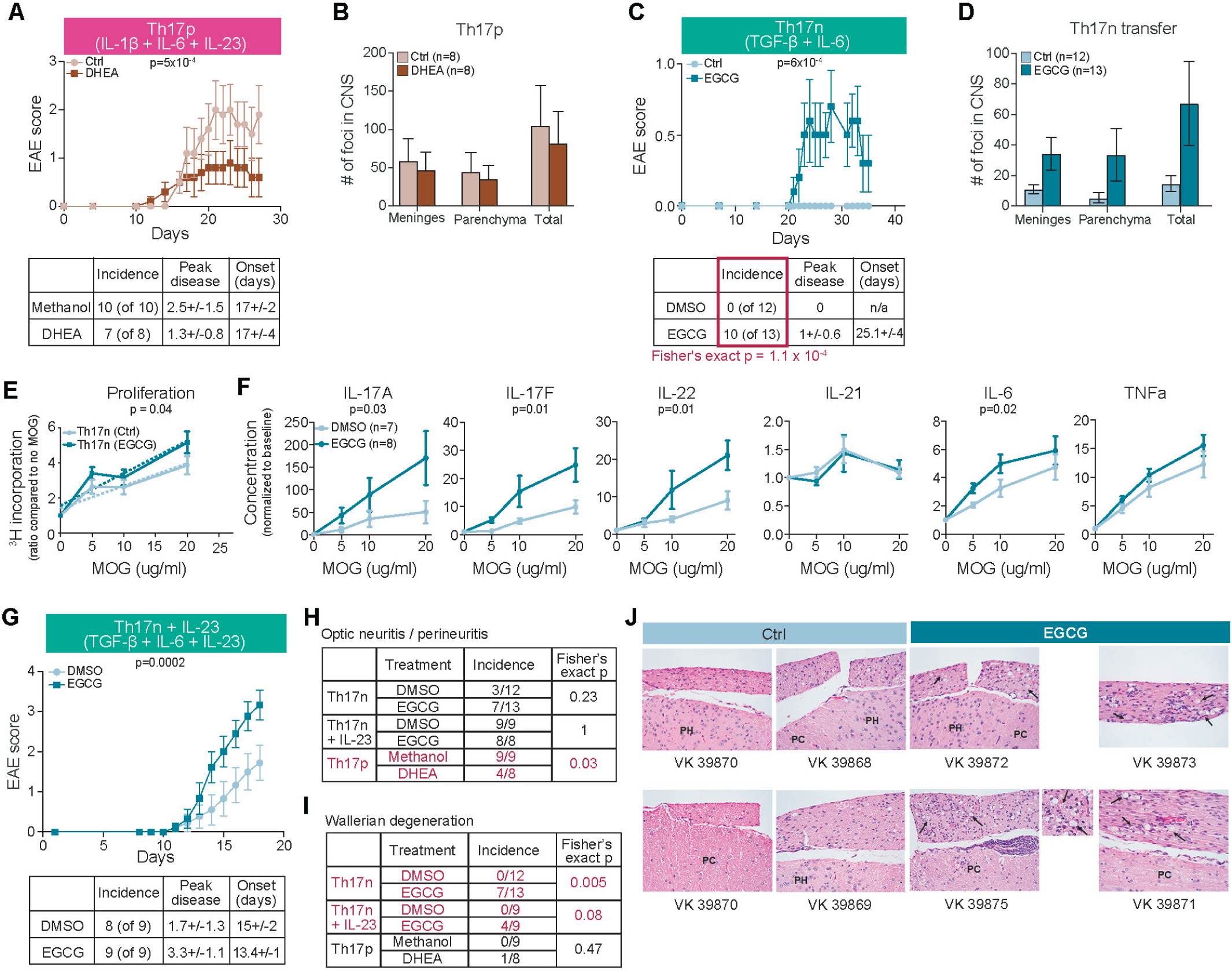
PGAM inhibition suppresses a regulatory Th17n program. (A) UMAP of single-set dataset; (B) Density plot of metabolic activity across cells. Metabolic activity was defined as the portion of the transcriptome consisting of RNA coding for metabolic enzymes. (C) Each cell was scored with the pro-inflammatory, pro-regulatory, or the difference between the pro-inflammatory and pro-regulatory signatures. Signatures were derived from transcriptomic modules we have previously described **(Methods)**. The difference between the signatures (rightmost boxes) is the overall pathogenicity score. (D) Genes with a significant (p < 0.001) interaction term in a linear model with cytokine stimulation and glucose concentration, and which were annotated as related to Th17 effector functions **(Methods)** are shown. The mean expression of each gene was z-scaled across the four conditions. Genes colored in red are discussed in the main text pertaining to the panel. (E) Same UMAP as in (A) colored by Leiden clusters. Each quadrant shows only cells belonging to one of the four conditions in color. (F) Genes that were differentially expressed between the clusters and annotated as related to Th17 effector functions **(Methods)** are shown. (G) The pathogenicity score of each cell was computed. Cells in the N1 cluster scored significantly lower than cells in the other clusters. (H) An EGCG signature, measuring the similarity of a Th17n cell to an EGCG-treated Th17n cell, was computed **(Methods)**. Cells in the N1 cluster score significantly lower than cells in the other clusters. (I-J) Gene signatures based on the defining markers for each Leiden cluster were computed **(Methods)**. GSEA plots showing the running sum statistic^34^ are shown. Horizontal dashed line denote the height at which each curve attains its maximum or minimum, corresponding to the GSEA enrichment score. Bars at the bottom correspond to indices of genes belonging to one of the three sets.

For a more granular analysis of the effect of glucose concentration, we applied a multivariate model for differential expression that accounted for the glucose level, the differentiation condition, and their interaction term. The effects of the differentiation protocols were largely consistent with the previous literature, capturing genes with known pro-inflammatory or pro-regulatory functions. However, we detected transcriptional responses to the increase in glucose that were specific for each differentiation condition. In the pathogenic (Th17p) condition, we found transcription factors, cytokines, chemokines, and surface proteins such as TBX21 (T-bet), CCL5, IL23 receptor, and IL22 whose expression increases in the high glucose conditions, whereas CSF2 (GM-CSF), a key regulator of Th17 cell pathogenicity and EAE, as well as GZMB, are preferentially expressed under the low glucose condition (**Figure 3D**). In the non-pathogenic (Th17n) condition, FOXP3, CTLA4, IL2RA (CD25), and TSC22D3 (which we previously identified as a Th17 regulator^21^) were all sensitive to glucose concentration. Thus, some drivers of Th17 cell function are sensitive to nutrient availability, and their perturbation may have different consequences in different tissue environments.

To further probe the diversity between cells (even in each condition), we partitioned our extended scRNAseq data set into clusters **(Methods)**. We identified three Th17n programs (N1-N3) and four Th17p programs (P1-P4) (**Figure 3E**). The programs were characterized by distinct Th17 effector markers (**Figure 3F; Table S3**). Within Th17p cells, clusters P1 and P2 were abundant in the presence of low glucose and had high expression of different effector molecules (GM-CSF in P1 vs. GZMB in P2). Cluster P3 was found predominantly under high glucose conditions and had high expression of TXNIP, which is known to be induced by glucose, and crucial for NLRP3 inflammasome activation. Finally, cluster P4 was characterized by the highest expression of IL22 and IL23R. In Th17n cells, cluster N2 was mostly restricted to low-glucose conditions and had high expression of LGALS3 and IL23a. Cluster N3 was found predominantly in high glucose conditions and had the highest expression of TNF and IL21. Cluster N1, which had high expression of FOXP3 and SGK1, was the only cluster robustly detected under both glucose conditions.

Th17n cells belonging to cluster N1 were less pathogenic than other Th17n cells **(Figure 3G)**. We hypothesized that EGCG treatment may specifically suppress this program. To test this, we scored every cell in the extended dataset according to its expression of the EGCG-response signature (as in **Figure 2E**). We found that N1 cells scored significantly lower than cells of other clusters (i.e., their transcriptomic program is negatively associated with EGCG effects; **Figure 3H)**. GSEA analysis further confirmed this trend **(Methods)** – we find that genes that are up-regulated in N1 cells (compared to all other Th17n cells) were significantly enriched in genes downregulated by EGCG **(Figure 3I).** We did not observe similar trends for the other clusters. Similarly, there was no strong association between the effects of DHEA, and the transcriptional programs of the different clusters **(Figure 3J)**. Taken together, we find that EGCG treatment results in suppression of genes that characterize the N1 phenotype more potently than other Th17n programs, thereby suggesting that loss or reprogramming of the N1 subset may underlie the shift of Th17n cells towards a pro-inflammatory phenotype.

### PGAM inhibition exacerbates, whereas G6PD inhibition ameliorates, Th17-mediated neuroinflammation *in vivo*

To test the functional relevance of the transcriptome shifts induced by EGCG and DHEA *in vivo*, we induced EAE by adoptive transfer of Th17 cells treated with various inhibitors in vitro, thus restricting the effect of the inhibitors specifically to the disease-inducing Th17 cells. We generated Th17n and Th17p cells from naïve CD4+ T cells isolated from 2D2 TCR-transgenic mice, with specificity for MOG35-55 peptide, and transferred them into wild-type mice to induce EAE.

First, we evaluated the effect of DHEA-treated Th17 cells on the disease development. Transfer of 2D2 T cells differentiated in the presence of DHEA under Th17p conditions reduced the severity of peak disease in the recipient mice (**Figure 4A**). However, disease incidence was unchanged by DHEA, and the number of lesions in the CNS was not significantly different (**Figure 4B**). Additionally, there were no significant alterations in antigen-specific cytokine secretion in response to MOG peptide, except for an increase in IL2 in the DHEA-treated group (**Figure S3A**). This is consistent with the in vitro transcriptome analysis, which suggested that DHEA did not restrict the proinflammatory program (**Figure 2D**).

**Figure 4.**
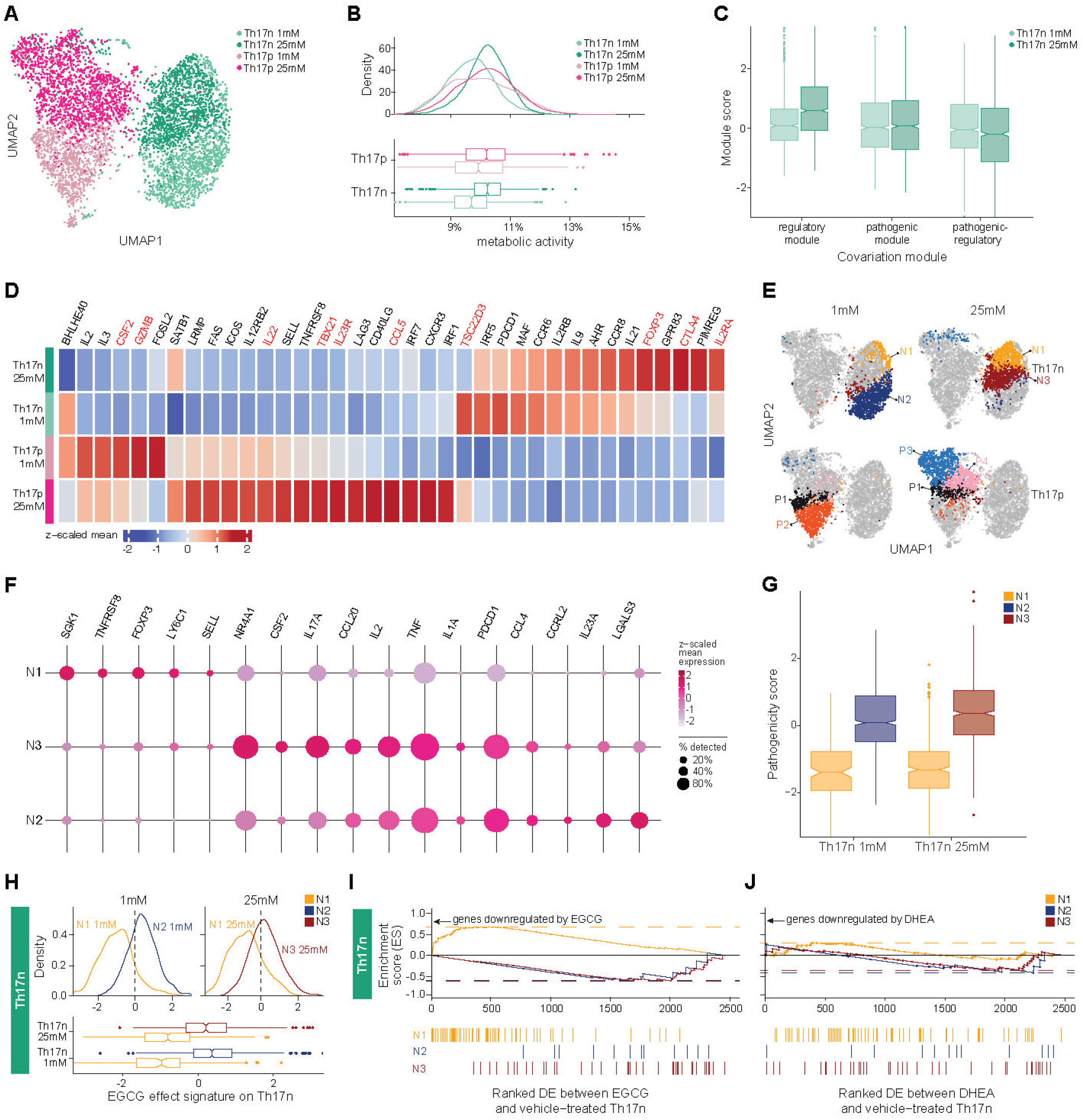
PGAM inhibition exacerbates, whereas G6PD inhibition ameliorates, Th17-mediated neuroinflammation *in vivo.* 2D2 TCR–transgenic Th17 cells were adoptively transferred after differentiation in vitro in the presence of an inhibitor or vehicle as indicated. (A, C, G) Clinical outcome of EAE; p-values test the null hypothesis of equal slopes of the EAE score curves in linear regression. (B, D) Histological score based on cell infiltrates in meninges and parenchyma of CNS; (E, F) Draining lymph node (cervical) from respective mice were isolated and pulsed with increasing dose of MOG_35-55_ peptide for 3 days and (E) subjected to thymidine incorporation assay; p-values test the null hypothesis of equal slopes for the two curves. by linear regression; or (F) measurement of cytokine secretion by Legendplex and flow cytometry. Concentrations were normalized through division by the respective response to no antigen control (H-I) Independent pathological report of CNS isolated from mice with EAE at endpoint (d35 for EGCG experiments; d28 for DHEA experiment); Optic nerves were not found in the histologic section from one animal in the EGCG+IL23 group. (J) Representative histology of spinal cord and spinal nerve roots. There is greater meningeal inflammation and Wallerian degeneration (digestion chambers, arrows) in posterior spinal nerve roots in EGCG vs. Control mice. PC, posterior column; PH, posterior horn. Individual mouse numbers are indicated. The smaller panel shows VK 39875 mouse section at higher magnification. All are H. & E., 40X objective. Three similar experiments were performed.

More interestingly, and in agreement with Compass predictions and our in-vitro analyses, EGCG-treated non-pathogenic Th17 (Th17n) cells induced EAE compared to the effect of vehicle-treated cells in recipient mice **(Figure 4C)**. Recipients of EGCG-treated Th17n cells had a significantly higher EAE incidence rate (10/12) compared with the control group (0/12, Fisher’s exact p = 1.1*10^-4^). Consistent with the clinical disease, histological analyses revealed an increased number of CNS lesions in both the meninges and the parenchyma of mice that received EGCG-treated Th17n cells (**Figure 4D**, p = 0.0093 by 2-way ANOVA). Analysis of MOG-specific responses showed that although there was only a small difference in antigen-specific T cell proliferation (**Figure 4E**), there was a significant increase in secretion of IL17, IL17F, IL22, and IL6 (**Figure 4F** and **Figure S3B**) in response to MOG antigen in cells isolated from the draining lymph node of the mice that had received EGCG-treated Th17n cells. As EGCG-treated non-pathogenic Th17n cells induced only mild EAE, we asked whether EGCG would further enhance the encephalitogenicity of Th17 cells in the presence of IL23, which enhances and stabilizes the Th17 phenotype ^23–26^. Indeed, transfer of Th17n cells differentiated in the presence of IL23 and EGCG significantly enhanced EAE disease severity, as compared with Th17n cells treated with IL23 and vehicle (**Figure 4G**).

In addition to CNS lesions, we performed additional histopathology analysis across all experiments. We found that while DHEA treatment in Th17p did not alter lesion load in meninges and parenchyma, it did not induce optic neuritis/perineuritis in host mice (**Figure 4H**). Conversely, EGCG treatment in Th17n (with or without IL23) had no such effect. Interestingly, mice transferred with EGCG-treated Th17 cells (Th17n or Th17n with IL23) were the only experimental group to produce Wallerian degeneration in proximal spinal nerve roots (**Figure 4I, J)**. Wallerian degeneration is not usually observed in the EAE model^27^, and indeed was not observed in our experiments even in EAE-inducing transfer of (untreated) Th17p or (untreated) Th17n+IL23. Overall, the distinct histopathology induced by Th17 cells treated with DHEA vis-à-vis EGCG further highlights the unique molecular programs underlying their biology.

In conclusion, we identified the glycolytic reaction PGAM as a key metabolic regulator suppressing Th17 pathogenicity in vivo. Inhibition of PGAM enhanced Th17-mediated disease, demonstrating the significance of this metabolic step. As a negative regulator of pathogenicity, PGAM adds complexity to the current view that the entire pathway of aerobic glycolysis was associated with a pro-inflammatory phenotype in Th17 cells.

## DISCUSSION

Glycolysis is critical for Th17 cell function and is considered a clinical target for Th17 cell-driven diseases. Here, we systematically interrogated glycolytic reactions in the context of Th17 cell pathogenicity, leveraging single-cell transcriptome data analyzed with the Compass algorithm. Our work interrogates specific glycolytic reactions and their roles in Th17 cell function and may inform target selection. Inhibition of G6PD and PKM restricted IL17 expression, whereas inhibition of PGAM promoted IL17 expression in Th17 cells, revealing complexity within the glycolysis pathway. Indeed, perturbation of G6PD and PGAM resulted in opposing effects on Th17 cell pathogenicity at the whole transcriptome level, consistent with the outcome of EAE induced by Th17 cell transfer.

Previous studies investigating glycolytic or related enzymes using complete genetic ablation approaches have all pointed to a critical role of glycolysis in immune cell maintenance^7–13^. This includes genetic deletion of PGAM in T cells, which attenuated both CD4 and CD8 T cell responses^28^. Other studies, however, highlighted the importance of studying not only complete ablation of genes but also their dose effects^29,30^ and their effects on normally-matured T cells in the circulation^31^, which may be more clinically relevant^32^. In this context, our results suggest that both chemical and genetic perturbation of PGAM can promote Th17 cell pathogenicity, placing PGAM in a unique position compared to other glycolytic enzymes. In line with our findings, blocking glycolysis using 2-DG was previously shown to promote Th17 cell effector function by activating cellular stress TGFb signaling networks^16^. PGAM is essential for TGFb signaling in cancer cells^33^ and may provide a molecular link between glycolysis and cellular stress. The effect of PGAM on Th17 cell pathogenicity may also be mediated through serine biosynthesis. Further studies are required to analyze the connection between PGAM, serine biosynthesis, cellular stress, and Th17 cell pathogenicity.

Our study also expanded the Th17 cell transcriptome atlas with data generated from Th17 cells cultured in both high (25mM) and low (1mM) glucose media. The different glucose conditions led to the emergence of different transcriptomic profiles, especially in the case of Th17p cells, highlighting the importance of the metabolic environment alongside the cytokine milieu in determining Th17 phenotypes. Similarities between low-glucose and DHEA-suppressed Th17p (e.g., upregulation of GZMB and downregulation of TBX21, IL22) suggest that DHEA works, in part, by mimicking glucose restriction.

We used the single-cell resolution to detect distinct transcriptomic programs, which likely correspond to distinct Th17 effector states, and occur in response to a specific combination of differentiation factors and glucose availability. Notably, one program of each differentiation condition (N1 and P1) occurred under both the high- and low-glucose conditions, whereas the other programs occurred only under a certain combination of glucose abundance and differentiating cytokines (**Figure 3F**). The N1 and P1 programs straddled the low and high glucose conditions but mostly belonged to the high glucose condition. They may represent cells that, even in the glucose-poor conditions, were able to sequester more glucose, or alternatively, a subset of cells transcriptomically primed to take better advantage of nutrients they may encounter even in a resource-poor environment. Intriguingly, the predominant effect of PGAM inhibition was on the N1 cluster that is induced in Th17n cells under both high and low glucose conditions. Compared to other Th17n programs, the N1 cluster showed the lowest pathogenicity score (**Figure 3G**), highlighting a role for PGAM in maintaining a niche pro-regulatory state of Th17n cells.

Cellular metabolism exerts major effects on immune cell function. With the increasing number of single-cell transcriptome datasets generated in human and preclinical models, our study provides further support to the application of Compass across data sets generated using different technology platforms, enabling systematic analysis of metabolic reactions and their connections to cellular function at single-cell resolution. Here, the use of Compass allowed us to detect a unique role for PGAM in determining the pathogenic state of Th17 cells in the context of autoimmunity, which sets it apart from other enzymes belonging to the glycolysis pathway. Our results suggest that reactions belonging to the same metabolic pathway can still modulate immune function in different and even, in some cases, in opposing ways.

## AUTHOR CONTRIBUTIONS

CW and AW conceived and designed the study. VKK and NY supervised the study. AW, DD, and NY designed and performed computational analyses. CW, JF, designed, performed, and analyzed experiments, with significant contributions by YZ and SZ. KP, and CC performed the LC/MS metabolomics analysis. RS performed the histological analysis. CW, AW, NY, and VKK wrote the manuscript with contributions from all authors. All authors read and approved the final manuscript.

## ACKNOWLEDGMENTS

We thank Hanna Vega for scientific illustrations and Mary Collins for critical editing of the manuscript. CW is supported by the Canadian Institute of Health Research (CIHR PG498150) and a Career Transitional Fellowship from the National Multiple Sclerosis Society. NY and AW were supported by the Chan Zuckerberg Biohub and by a National Institute of Mental Health (NIMH) grant NIH5U19MH114821. JF was supported by a Max Kade fellowship awarded by the Austrian Academy of Science (ÖAW). This work was supported by grants from the National Institutes of Health (RO1NS30843, R01AI144166, R01NS045937, P01AI073748, P01AI039671, P01AI056299) awarded to VKK.

## SUPPLEMENTARY FIGURE LEGENDS

**Figure S1.**
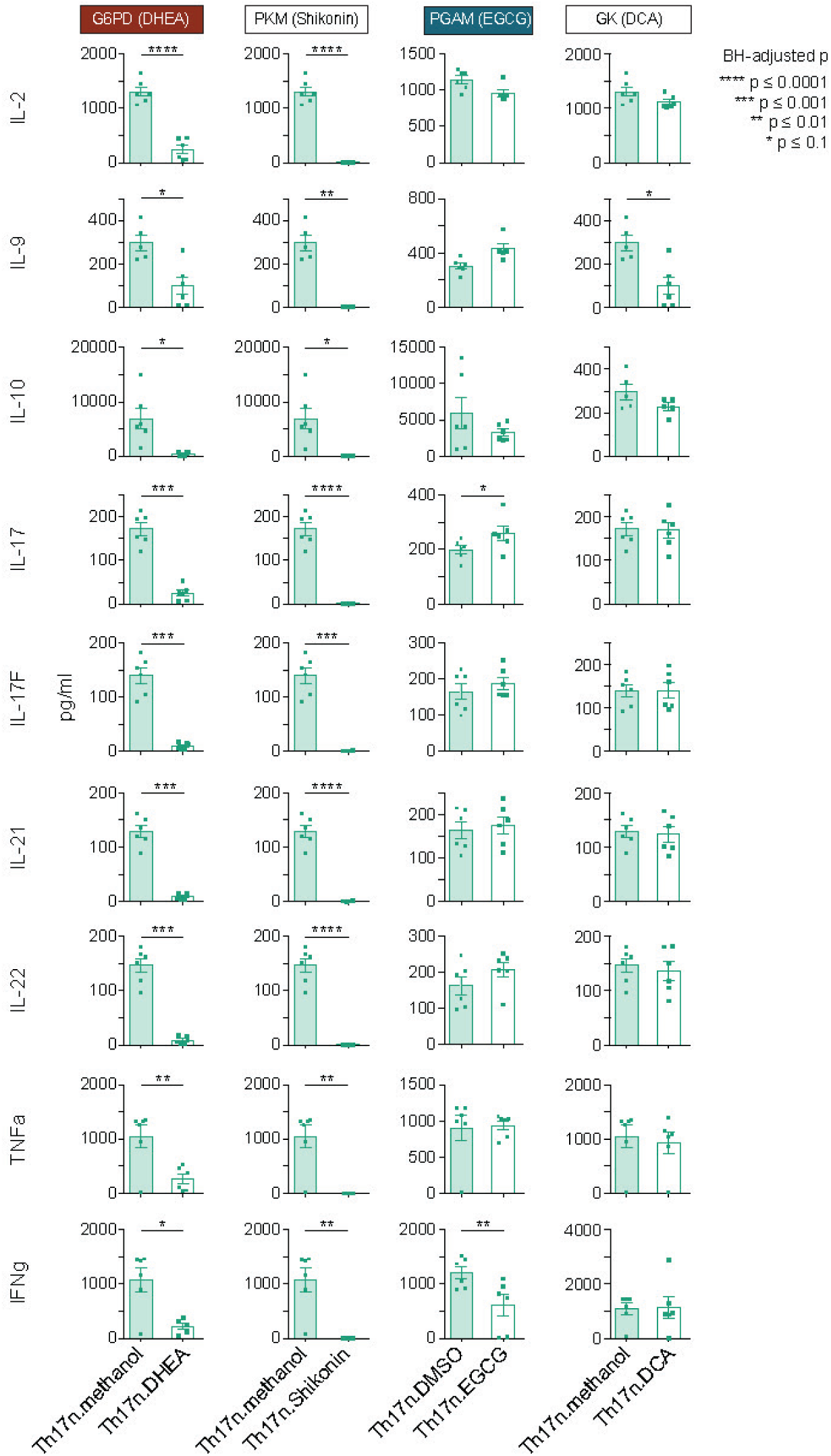
(A) Supernatant from Th17 cell cultures performed for main Figure 1d are harvested for cytokine analysis using Legendplex.

**Figure S2.**
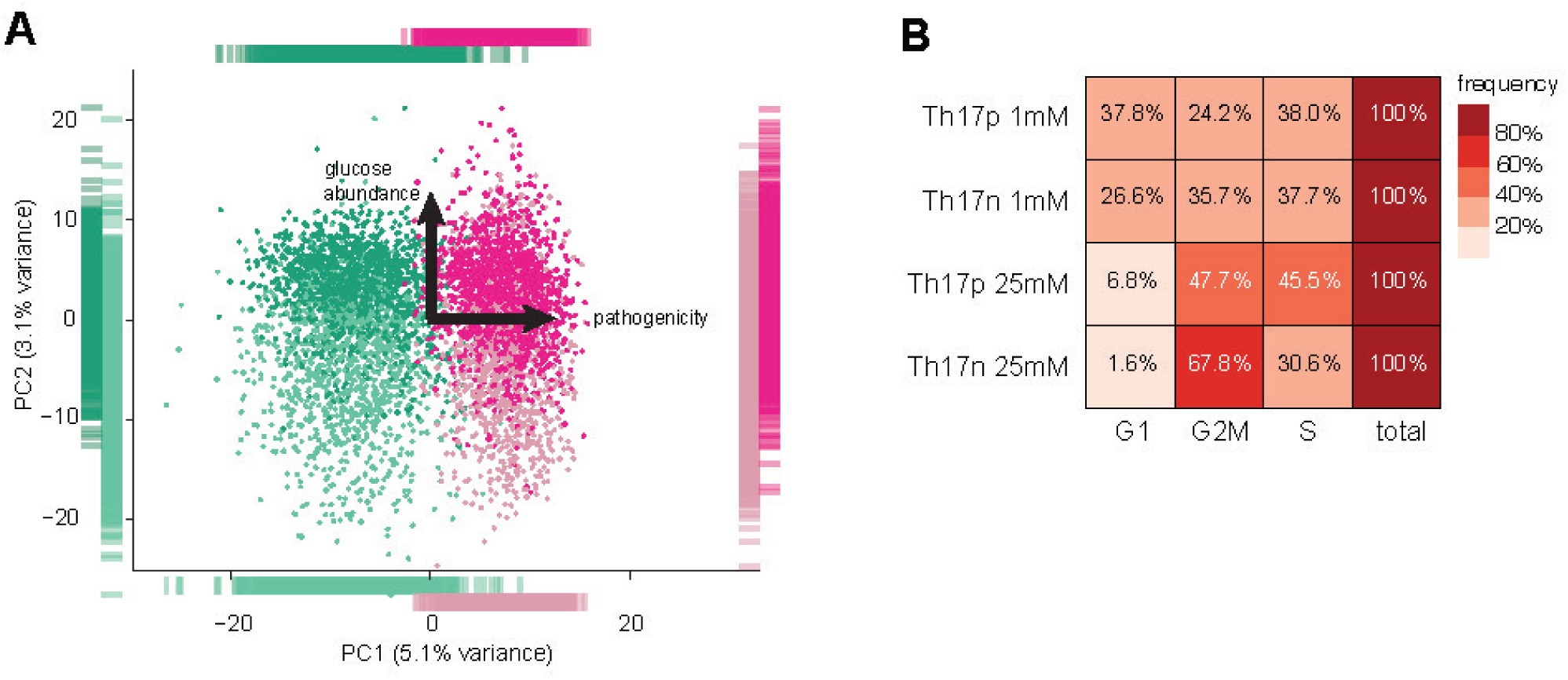
(A) PCA of the same data shown in Figure 3A. (B) Ratio of cells belonging to the different cell cycle phases. Cells were assigned into cell cycle phases with Scanpy’s `score_genes_cell_cycle` function^35^.

**Figure S3.**
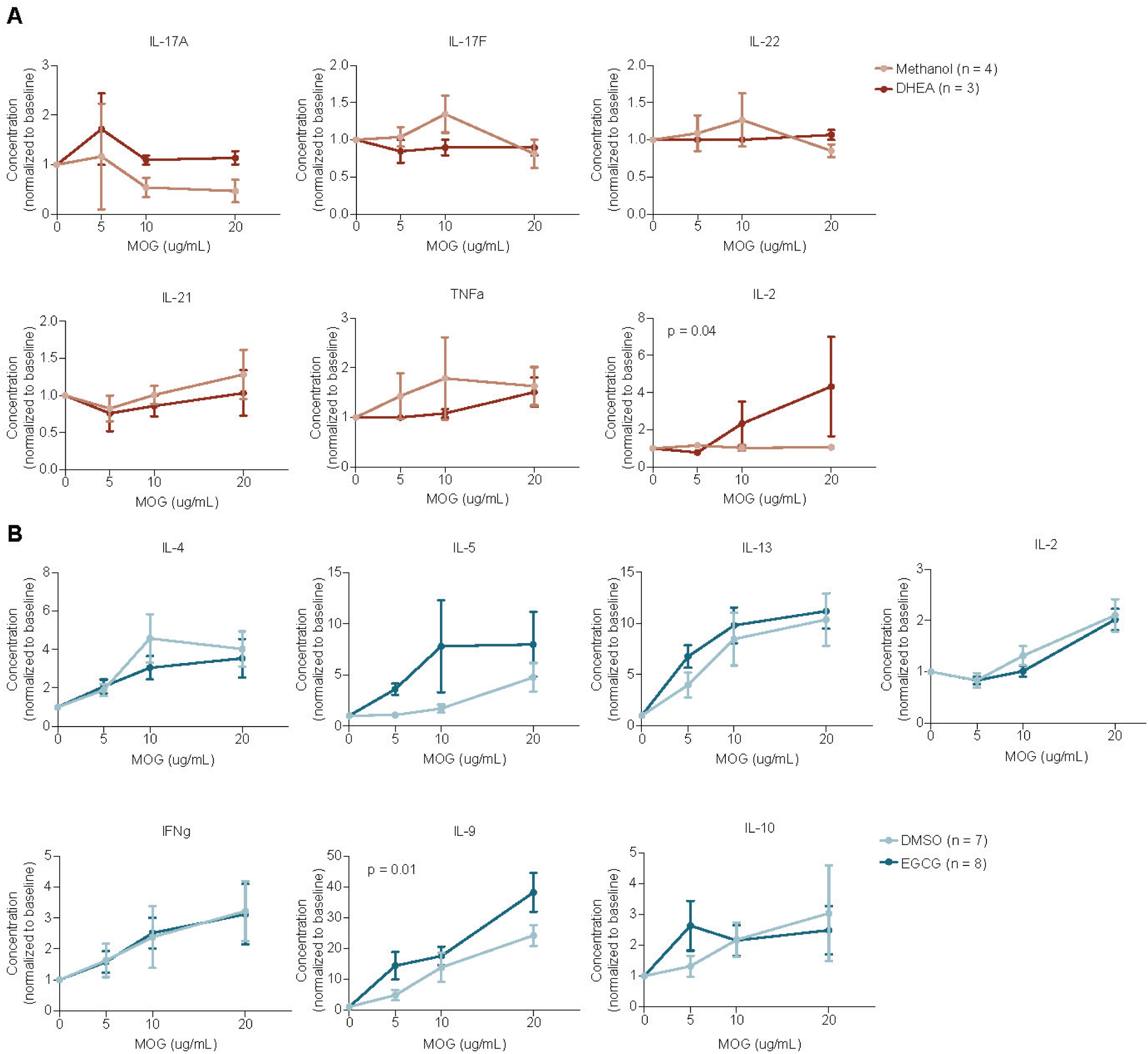
Cytokine secretion after three days of culture with increasing dose of MOG_35-55_ peptide from cells isolated from draining lymph node (cervical) of mice transferred with (A) methanol or DHEA treated Th17p cells as in Figure 4A or (B) DMSO or EGCG as in Figure 4C. Concentrations were normalized through division by the respective response to no antigen control.

## METHODS

### T cell differentiation culture

Naive CD4+CD44-CD62L+CD25-T cells were sorted using BD FACSAria sorter and activated with plate-bound anti-CD3 and anti-CD28 antibodies (both at 1mg/ml) in the presence of cytokines at a concentration of 0.5X10^6^ cells/ml. For Th17 differentiation: 2ng/ml of rhTGFb1, 25ng/ml rmIL-6, 20ng/ml rmIL-1b (all from Miltenyi Biotec) and 20ng/ml rmIL-23 (R & D systems) were used at various combinations as specified in figures. For differentiation experiments, cells were harvested at 68 hours for RNA analysis and 72-96h for flow cytometry analysis. Cells were re-stimulated with PMA/ionomycin for four hours in the presence of brefaldin and monensin before analysis for cytokines by intracellular cytokine staining. Cytokine concentrations in supernatants of in vitro cultures were analyzed by the Legendplex Mouse Th Cytokine Panel (13-plex) (BioLegend) according to the manufacturer’s instructions and analyzed on a FACS LSR II (BD Biosciences).

### Chemical Inhibitors

All chemical inhibitors were purchased from Sigma except EGCG (Selleck Chemicals) and tested in a wide range of dose (20nM-200uM) on Th17 cells. The lowest dose that resulted in minimal impact on cell viability is used for functional evaluation: EGCG, the inhibitor for PGAM^36^, was used at 20-50uM; DHEA, inhibitor for G6PD^37^, was used at 50uM; DCA, inhibitor for GK^38,39^, was used at 40uM; and Shikonin, inhibitor for PKM2^40,41^, was used at 10uM.

### PGAM genetic perturbation

The initial CRISPR guide sequences were selected based on the exon structure of target genes and ranked by the repertoire of potential off-target sites to select designs that minimize the possibility of off-target cleavage. The guides were then cloned into CRISPR-Cas9 vectors (pck005-tdtomato, Addgene #85453) via golden-gate cloning as previously described^42^. The final guide sequence selected for PGAM1 was “TAAGTGCGCAGGCGCGACCG”. Control or PGAM Lentivirus was produced using 293T cells and used to infect Th17n cells differentiated *in vitro*. Spleen of Cas9-expressing mice (Jackson) were used as a source of T cells.

### Mice

C57BL/6 wildtype (WT) mice were obtained from Jackson Laboratory. WT 2D2 transgenic mice were bred in-house. All experiments were performed in accordance with the guidelines outlined by the Harvard Medical Area Standing Committee on Animals at the Harvard Medical School (Boston, MA).

### Experimental Autoimmune Encephalomyelitis (EAE)

For adoptive transfer EAE, naive T cells (CD4+CD44-CD62L+CD25-) were isolated from 2D2 TCR-transgenic mice and activated with anti-CD3 (1mg /ml) and anti-CD28 (1mg /ml) in the presence of differentiation cytokines for 68h. Cells were rested for 2 days and restimulated with plate-bound anti-CD3 (0.5mg/ml for pathogenic condition; 1mg /ml for non-pathogenic condition) and anti-CD28 (1mg /ml) for 2 days prior to transfer. Equal numbers (2 to 8 million) of cells were transferred per mouse intravenously. EAE is scored as previously published^27^.

### LC/MS metabolomics and carbon tracing

For carbon tracing experiments, Th17 cells were differentiated as described. Thereafter, at 96h, cells were washed and cultured in media supplemented with 8 mM [U-13C]-glucose for 15min. Samples were snap-frozen and extracted in 80% methanol. Two liquid chromatography tandem mass spectrometry (LC-MS) methods were used to measure fatty acids and lipids in cell extracts.

### RNA-seq processing

Cultured Th17 cells from various conditions were processed for either population RNA-seq (chemical inhibition) or scRNA-seq (25mM vs. 1mM glucose) using the 10x genomics platform (5’ GEM). Samples from 2-3 mice per group were processed with CellRanger. Preliminary analysis for quality and exclusion of low-quality droplets were performed with Scanpy^35^. Low-quality droplets were excluded. Cells were assigned into cell cycle phases with Scanpy’s `score_genes_cell_cycle` function^35^, with gene sets corresponding to cell cycle phases as described in Scanpy’s documentation. Highly variable genes (HVGs) were detected with Scanpy’s `highly_variable_genes` function, setting flavor=”seurat”, and separating to batches by animal, ultimately selecting 1,429 HVGs.

The high-dimensional HVG space was reduced to a 30-dimension latent space with scvi-tools^43^, using a negative-binomial generative model, with two discrete nuisance covariates per cell: animal and cell cycle phase. The cell cycle phase was regressed out after we observed it dominated the structure of the data. We then used Scanpy to compute a shared nearest neighbor (SNN) graph and Leiden clustering on top of the SNN graph. We identified clusters corresponding to low-quality or undifferentiated cells, excluded them, and repeated the analysis pipeline described above to generate the final latent scVI space. The final dataset consisted of 5,192 cells.

### Glucose- and cytokine-specific Th17 states

We used Scanpy to generate an SNN graph based on distances the final latent space. We computed Leiden clusters for the SNN graph, setting the Leiden’s resolution parameter to 0.8, which resulted in 10 clusters. We excluded three clusters that together contained 110 cells. The remaining seven clusters (N1-N3, P1-P4) were either a) specific to a cell type and detected in both glucose concentration (N1, P1); or b) specific to a cell type and glucose concentration (N2-N3, P2-P4). We defined a cluster as specific to a cell type or cell type and glucose combination if it had no more than 100 cells in each of the other conditions.

### Transcriptomic signatures

Transcriptomic signatures were computed as previously described^17^. Given a list of genes, we compute a signature score by z-scaling the log-scale expression of each gene across cells and taking an average. If the list contains both genes that are positively associated and genes that are negatively associated to the signature, we multiply the z-scaled expression by +1 or -1, respectively, prior to taking the average. Only highly variable genes (HVGs) are considered for the purpose of signature computation.

The pathogenicity signature was based on the pro-inflammatory and pro-regulatory transcriptional modules we have previously identified^22^ (**Table S1;** see Figure 4b in that publication). The pathogenicity score was computed by subtracting the pro-inflammatory module score from the pro-regulatory module score. Out of the 116 genes in the pro-inflammatory module, only 63 that were also HVG in our dataset were considered for signature computation. Similarly, out of 68 genes in the pro-regulatory module, only 30 HVGs were considered.

EGCG effect signature was defined as the set of differentially expressed genes between EGCG- and vehicle-treated cells in bulk RNA-seq, with a sign of +1 or -1 for up- and downregulated genes, respectively. Differentially expressed genes were computed with limma, with thresholds of abs(logFC) >= log2(1.5) and BH-adjusted p-value <= 0.05 **(Table S2)**. DHEA effect signature was similarly computed.

### Compass analysis

To mitigate 10x count sparsity, the input to Compass was normalized counts were imputed by the scVI model with the `SCVI.get_normalized_expression` function. Compass was run as previously described^17^, with the following adjustments:

a. The imputation with scVI already introduced information-sharing between cells, and we therefore disabled further information sharing by setting the lambda parameter to 0.
b. Meta-reactions were not used to focus on reaction-level results.

### Gene sets and gene-set enrichment analysis (GSEA)

We defined a set of genes that are broadly related to Th17 effector function by manually curating relevant literature^19–22,44–52^. Genes belonging to relevant families, such as cytokine,cytokine receptors, interferon response were added to the set.

Gene sets associated with the transcriptional modules N1-N3 **(Table S3)** were defined by selecting genes that were upregulated in one of the modules compared to all other Th17n cells in the dataset, as decided by Seurat’s `FindAllMarkers` function^53^. Threshold were set at abs(avg. logFC) > 1) and BH-adjusted p-value < 1e-5. GSEA statistic^34^ was computed with the `fgsea` package.

## REFERENCES

1. Dang, E.V., Barbi, J., Yang, H.-Y., Jinasena, D., Yu, H., Zheng, Y., Bordman, Z., Fu, J., Kim, Y., Yen, H.-R., et al. (2011). Control of T(H)17/T(reg) balance by hypoxia-inducible factor 1. Cell 146, 772–784.

2. Damasceno, L.E.A., Prado, D.S., Veras, F.P., Fonseca, M.M., Toller-Kawahisa, J.E., Rosa, M.H., Públio, G.A., Martins, T.V., Ramalho, F.S., Waisman, A., et al. (2020). PKM2 promotes Th17 cell differentiation and autoimmune inflammation by fine-tuning STAT3 activation. J. Exp. Med. 217. 10.1084/jem.20190613.

3. Hochrein, S.M., Wu, H., Eckstein, M., Arrigoni, L., Herman, J.S., Schumacher, F., Gerecke, C., Rosenfeldt, M., Grün, D., Kleuser, B., et al. (2022). The glucose transporter GLUT3 controls T helper 17 cell responses through glycolytic-epigenetic reprogramming. Cell Metab. 34, 516–532.e11.

4. Gerriets, V.A., Kishton, R.J., Nichols, A.G., Macintyre, A.N., Inoue, M., Ilkayeva, O., Winter, P.S., Liu, X., Priyadharshini, B., Slawinska, M.E., et al. (2015). Metabolic programming and PDHK1 control CD4+ T cell subsets and inflammation. J. Clin. Invest. 125, 194–207.

5. Xu, K., Yin, N., Peng, M., Stamatiades, E.G., Chhangawala, S., Shyu, A., Li, P., Zhang, X., Do, M.H., Capistrano, K.J., et al. (2021). Glycolytic ATP fuels phosphoinositide 3-kinase signaling to support effector T helper 17 cell responses. Immunity 54, 976–987.e7.

6. Wu, L., Hollinshead, K.E.R., Hao, Y., Au, C., Kroehling, L., Ng, C., Lin, W.-Y., Li, D., Silva, H.M., Shin, J., et al. (2020). Niche-Selective Inhibition of Pathogenic Th17 Cells by Targeting Metabolic Redundancy. Cell 182, 641–654.e20.

7. Pearce, E.L., Poffenberger, M.C., Chang, C.-H., and Jones, R.G. (2013). Fueling Immunity: Insights into Metabolism and Lymphocyte Function. Science 342. 10.1126/science.1242454.

8. Pearce, E.J., and Everts, B. (2015). Dendritic cell metabolism. Nat. Rev. Immunol. 15, 18–29.

9. MacIver, N.J., Michalek, R.D., and Rathmell, J.C. (2013). Metabolic Regulation of T Lymphocytes. Annu. Rev. Immunol. 31, 259–283.

10. O’Neill, L.A.J., and Pearce, E.J. (2016). Immunometabolism governs dendritic cell and macrophage function. J. Exp. Med. 213, 15–23.

11. Geltink, R.I.K., Kyle, R.L., and Pearce, E.L. (2018). Unraveling the Complex Interplay Between T Cell Metabolism and Function. Annu. Rev. Immunol. 36, 461–488.

12. O’Neill, L.A.J., Kishton, R.J., and Rathmell, J. (2016). A guide to immunometabolism for immunologists. Nat. Rev. Immunol. 16, 553–565.

13. O’Brien, K.L., and Finlay, D.K. (2019). Immunometabolism and natural killer cell responses. Nat. Rev. Immunol. 19, 282–290.

14. Van den Bossche, J., O’Neill, L.A., and Menon, D. (2017). Macrophage Immunometabolism: Where Are We (Going)? Trends Immunol. 38, 395–406.

15. Newton, R., Priyadharshini, B., and Turka, L.A. (2016). Immunometabolism of regulatory T cells. Nat. Immunol. 17, 618–625.

16. Brucklacher-Waldert, V., Ferreira, C., Stebegg, M., Fesneau, O., Innocentin, S., Marie, J.C., and Veldhoen, M. (2017). Cellular Stress in the Context of an Inflammatory Environment Supports TGF-β-Independent T Helper-17 Differentiation. Cell Rep. 19, 2357–2370.

17. Wagner, A., Wang, C., Fessler, J., DeTomaso, D., Avila-Pacheco, J., Kaminski, J., Zaghouani, S., Christian, E., Thakore, P., Schellhaass, B., et al. (2021). Metabolic modeling of single Th17 cells reveals regulators of autoimmunity. Cell 184, 4168–4185.e21.

18. Pan, Y., Yang, W., Tang, B., Wang, X., Zhang, Q., Li, W., and Li, L. (2023). The protective and pathogenic role of Th17 cell plasticity and function in the tumor microenvironment. Front. Immunol. 14. 10.3389/fimmu.2023.1192303.

19. Lee, Y., Awasthi, A., Yosef, N., Quintana, F.J., Xiao, S., Peters, A., Wu, C., Kleinewietfeld, M., Kunder, S., Hafler, D.A., et al. (2012). Induction and molecular signature of pathogenic TH17 cells. Nat. Immunol. 13, 991–999.

20. Wang, C., Yosef, N., Gaublomme, J., Wu, C., Lee, Y., Clish, C.B., Kaminski, J., Xiao, S., Horste, G.M.Z., Pawlak, M., et al. (2015). CD5L/AIM Regulates Lipid Biosynthesis and Restrains Th17 Cell Pathogenicity. Cell 163, 1413–1427.

21. Yosef, N., Shalek, A.K., Gaublomme, J.T., Jin, H., Lee, Y., Awasthi, A., Wu, C., Karwacz, K., Xiao, S., Jorgolli, M., et al. (2013). Dynamic regulatory network controlling TH17 cell differentiation. Nature 496, 461–468.

22. Gaublomme, J.T., Yosef, N., Lee, Y., Gertner, R.S., Yang, L.V., Wu, C., Pandolfi, P.P., Mak, T., Satija, R., Shalek, A.K., et al. (2015). Single-Cell Genomics Unveils Critical Regulators of Th17 Cell Pathogenicity. Cell 163, 1400–1412.

23. Awasthi, A., Riol-Blanco, L., Jäger, A., Korn, T., Pot, C., Galileos, G., Bettelli, E., Kuchroo, V.K., and Oukka, M. (2009). Cutting edge: IL-23 receptor gfp reporter mice reveal distinct populations of IL-17-producing cells. J. Immunol. 182, 5904–5908.

24. McGeachy, M.J., Chen, Y., Tato, C.M., Laurence, A., Joyce-Shaikh, B., Blumenschein, W.M., McClanahan, T.K., O’Shea, J.J., and Cua, D.J. (2009). The interleukin 23 receptor is essential for the terminal differentiation of interleukin 17-producing effector T helper cells in vivo. Nat. Immunol. 10, 314–324.

25. Zhou, L., Ivanov, I.I., Spolski, R., Min, R., Shenderov, K., Egawa, T., Levy, D.E., Leonard, W.J., and Littman, D.R. (2007). IL-6 programs T(H)-17 cell differentiation by promoting sequential engagement of the IL-21 and IL-23 pathways. Nat. Immunol. 8, 967–974.

26. Aggarwal, S., Ghilardi, N., Xie, M.-H., de Sauvage, F.J., and Gurney, A.L. (2003). Interleukin-23 promotes a distinct CD4 T cell activation state characterized by the production of interleukin-17. J. Biol. Chem. 278, 1910–1914.

27. Jäger, A., Dardalhon, V., Sobel, R.A., Bettelli, E., and Kuchroo, V.K. (2009). Th1, Th17, and Th9 effector cells induce experimental autoimmune encephalomyelitis with different pathological phenotypes. J. Immunol. 183, 7169–7177.

28. Toriyama, K., Kuwahara, M., Kondoh, H., Mikawa, T., Takemori, N., Konishi, A., Yorozuya, T., Yamada, T., Soga, T., Shiraishi, A., et al. (2020). T cell-specific deletion of Pgam1 reveals a critical role for glycolysis in T cell responses. Commun Biol 3, 394.

29. Chauhan, S.K., Saban, D.R., Lee, H.K., and Dana, R. (2009). Levels of Foxp3 in regulatory T cells reflect their functional status in transplantation. J. Immunol. 182, 148–153.

30. Gubin, M.M., Techasintana, P., Magee, J.D., Dahm, G.M., Calaluce, R., Martindale, J.L., Whitney, M.S., Franklin, C.L., Besch-Williford, C., Hollingsworth, J.W., et al. (2014). Conditional knockout of the RNA-binding protein HuR in CD4^+^ T cells reveals a gene dosage effect on cytokine production. Mol. Med. 20, 93–108.

31. Locke, F.L., Zha, Y.-Y., Zheng, Y., Driessens, G., and Gajewski, T.F. (2013). Conditional deletion of PTEN in peripheral T cells augments TCR-mediated activation but does not abrogate CD28 dependency or prevent anergy induction. J. Immunol. 191, 1677–1685.

32. Kuehn, H.S., Ouyang, W., Lo, B., Deenick, E.K., Niemela, J.E., Avery, D.T., Schickel, J.-N., Tran, D.Q., Stoddard, J., Zhang, Y., et al. (2014). Immune dysregulation in human subjects with heterozygous germline mutations in CTLA4. Science 345, 1623–1627.

33. Huang, K., Liang, Q., Zhou, Y., Jiang, L.-L., Gu, W.-M., Luo, M.-Y., Tang, Y.-B., Wang, Y., Lu, W., Huang, M., et al. (2019). A Novel Allosteric Inhibitor of Phosphoglycerate Mutase 1 Suppresses Growth and Metastasis of Non-Small-Cell Lung Cancer. Cell Metab. 30, 1107–1119.e8.

34. Subramanian, A., Tamayo, P., Mootha, V.K., Mukherjee, S., Ebert, B.L., Gillette, M.A., Paulovich, A., Pomeroy, S.L., Golub, T.R., Lander, E.S., et al. (2005). Gene set enrichment analysis: A knowledge-based approach for interpreting genome-wide expression profiles. Proc. Natl. Acad. Sci. U. S. A. 102, 15545–15550.

35. Wolf, F.A., Angerer, P., and Theis, F.J. (2018). SCANPY: large-scale single-cell gene expression data analysis. Genome Biol. 19, 15.

36. Li, X., Tang, S., Wang, Q.-Q., Leung, E.L.-H., Jin, H., Huang, Y., Liu, J., Geng, M., Huang, M., Yuan, S., et al. (2017). Identification of Epigallocatechin-3-Gallate as an Inhibitor of Phosphoglycerate Mutase 1. Front. Pharmacol. 8, 325.

37. Schwartz, A.G., and Pashko, L.L. (2004). Dehydroepiandrosterone, glucose-6-phosphate dehydrogenase, and longevity. Ageing Res. Rev. 3, 171–187.

38. Westergaard, N., Madsen, P., and Lundgren, K. (1998). Characterization of glycerol uptake and glycerol kinase activity in rat hepatocytes cultured under different hormonal conditions. Biochim. Biophys. Acta 1402, 261–268.

39. Tisdale, M.J., and Threadgill, M.D. (1984). (+/-)2,3-Dihydroxypropyl dichloroacetate, an inhibitor of glycerol kinase. Cancer Biochem. Biophys. 7, 253–259.

40. Zhao, X., Zhu, Y., Hu, J., Jiang, L., Li, L., Jia, S., and Zen, K. (2018). Shikonin Inhibits Tumor Growth in Mice by Suppressing Pyruvate Kinase M2-mediated Aerobic Glycolysis. Sci. Rep. 8, 14517.

41. Chen, J., Xie, J., Jiang, Z., Wang, B., Wang, Y., and Hu, X. (2011). Shikonin and its analogs inhibit cancer cell glycolysis by targeting tumor pyruvate kinase-M2. Oncogene 30, 4297– 4306.

42. Cong, L., Ran, F.A., Cox, D., Lin, S., Barretto, R., Habib, N., Hsu, P.D., Wu, X., Jiang, W., Marraffini, L.A., et al. (2013). Multiplex genome engineering using CRISPR/Cas systems. Science 339, 819–823.

43. Gayoso, A., Lopez, R., Xing, G., Boyeau, P., Valiollah Pour Amiri, V., Hong, J., Wu, K., Jayasuriya, M., Mehlman, E., Langevin, M., et al. (2022). A Python library for probabilistic analysis of single-cell omics data. Nat. Biotechnol. 10.1038/s41587-021-01206-w.

44. Ciofani, M., Madar, A., Galan, C., Sellars, M., Mace, K., Pauli, F., Agarwal, A., Huang, W., Parkhurst, C.N., Muratet, M., et al. (2012). A validated regulatory network for Th17 cell specification. Cell 151, 289–303.

45. Wherry, E.J., and Kurachi, M. (2015). Molecular and cellular insights into T cell exhaustion. Nat. Rev. Immunol. 15, 486–499.

46. Xu, Z.-S., Zhang, H.-X., Li, W.-W., Ran, Y., Liu, T.-T., Xiong, M.-G., Li, Q.-L., Wang, S.-Y., Wu, M., Shu, H.-B., et al. (2019). FAM64A positively regulates STAT3 activity to promote Th17 differentiation and colitis-associated carcinogenesis. Proc. Natl. Acad. Sci. U. S. A. 116, 10447–10452.

47. Suzuki, A.S., Yagi, R., Kimura, M.Y., Iwamura, C., Shinoda, K., Onodera, A., Hirahara, K., Tumes, D.J., Koyama-Nasu, R., Iismaa, S.E., et al. (2020). Essential Role for CD30-Transglutaminase 2 Axis in Memory Th1 and Th17 Cell Generation. Front. Immunol. 11, 1536.

48. Yasuda, K., Kitagawa, Y., Kawakami, R., Isaka, Y., Watanabe, H., Kondoh, G., Kohwi-Shigematsu, T., Sakaguchi, S., and Hirota, K. (2019). Satb1 regulates the effector program of encephalitogenic tissue Th17 cells in chronic inflammation. Nat. Commun. 10, 549.

49. Lin, C.-C., Bradstreet, T.R., Schwarzkopf, E.A., Sim, J., Carrero, J.A., Chou, C., Cook, L.E., Egawa, T., Taneja, R., Murphy, T.L., et al. (2014). Bhlhe40 controls cytokine production by T cells and is essential for pathogenicity in autoimmune neuroinflammation. Nat. Commun. 5, 3551.

50. Wu, B., Zhang, S., Guo, Z., Bi, Y., Zhou, M., Li, P., Seyedsadr, M., Xu, X., Li, J.-L., Markovic-Plese, S., et al. (2021). The TGF-β superfamily cytokine Activin-A is induced during autoimmune neuroinflammation and drives pathogenic Th17 cell differentiation. Immunity 0. 10.1016/j.immuni.2020.12.010.

51. Deng, J., Pan, T., Liu, Z., McCarthy, C., Vicencio, J.M., Cao, L., Alfano, G., Suwaidan, A.A., Yin, M., Beatson, R., et al. (2023). The role of TXNIP in cancer: a fine balance between redox, metabolic, and immunological tumor control. Br. J. Cancer. 10.1038/s41416-023-02442-4.

52. Schnell, A., Huang, L., Singer, M., Singaraju, A., Barilla, R.M., Regan, B.M.L., Bollhagen, A., Thakore, P.I., Dionne, D., Delorey, T.M., et al. (2021). Stem-like intestinal Th17 cells give rise to pathogenic effector T cells during autoimmunity. Cell 0. 10.1016/j.cell.2021.11.018.

53. Hao, Y., Stuart, T., Kowalski, M.H., Choudhary, S., Hoffman, P., Hartman, A., Srivastava, A., Molla, G., Madad, S., Fernandez-Granda, C., et al. (2024). Dictionary learning for integrative, multimodal and scalable single-cell analysis. Nat. Biotechnol. 42, 293–304.

